# A Biomaterial Screening Approach Reveals Microenvironmental Mechanisms of Drug Resistance

**DOI:** 10.1101/168039

**Authors:** Alyssa D. Schwartz, Lauren E. Barney, Lauren E. Jansen, Thuy V. Nguyen, Christopher L. Hall, Aaron S. Meyer, Shelly R. Peyton

## Abstract

**TOC Figure:** Drug response screening, gene expression, and kinome signaling were combined across biomaterial platforms to combat adaptive resistance to sorafenib.

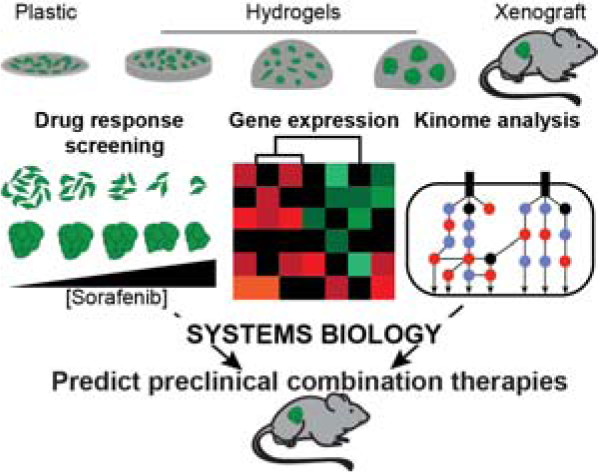

**Insight Box:** We combined biomaterial platforms, drug screening, and systems biology to identify mechanisms of extracellular matrix-mediated adaptive resistance to RTK-targeted cancer therapies. Drug response was significantly varied across biomaterials with altered stiffness, dimensionality, and cell-cell contacts, and kinome reprogramming was responsible for these differences in drug sensitivity. Screening across many platforms and applying a systems biology analysis were necessary to identify MEK phosphorylation as the key factor associated with variation in drug response. This method uncovered the combination therapy of sorafenib with a MEK inhibitor, which decreased viability on and within biomaterials *in vitro*, but was not captured by screening on tissue culture plastic alone. This combination therapy also reduced tumor burden *in vivo,* and revealed a promising approach for combating adaptive drug resistance.

**Abstract:** Traditional drug screening methods lack features of the tumor microenvironment that contribute to resistance. Most studies examine cell response in a single biomaterial platform in depth, leaving a gap in understanding how extracellular signals such as stiffness, dimensionality, and cell-cell contacts act independently or are integrated within a cell to affect either drug sensitivity or resistance. This is critically important, as adaptive resistance is mediated, at least in part, by the extracellular matrix (ECM) of the tumor microenvironment. We developed an approach to screen drug responses in cells cultured on 2D and in 3D biomaterial environments to explore how key features of ECM mediate drug response. This approach uncovered that cells on 2D hydrogels and spheroids encapsulated in 3D hydrogels were less responsive to receptor tyrosine kinase (RTK)-targeting drugs sorafenib and lapatinib, but not cytotoxic drugs, compared to single cells in hydrogels and cells on plastic. We found that transcriptomic differences between these *in vitro* models and tumor xenografts did not reveal mechanisms of ECM-mediated resistance to sorafenib. However, a systems biology analysis of phospho-kinome data uncovered that variation in MEK phosphorylation was associated with RTK-targeted drug resistance. Using sorafenib as a model drug, we found that co-administration with a MEK inhibitor decreased ECM-mediated resistance *in vitro* and reduced *in vivo* tumor burden compared to sorafenib alone. In sum, we provide a novel strategy for identifying and overcoming ECM-mediated resistance mechanisms by performing drug screening, phospho-kinome analysis, and systems biology across multiple biomaterial environments.

## Introduction

Adaptive resistance poses a significant challenge for targeted drugs as tumor cells can alter their signaling pathways to bypass the effect of therapeutics. This adaptive signaling occurs quickly and independently of acquired mutations, making the timing of therapeutics critical to sensitize tumor cells to drugs via rewiring from an oncogene-addicted state ^1^. For instance, triple-negative breast cancer cells treated with a MEK inhibitor undergo a rapid kinome reprogramming of many receptor tyrosine kinases (RTKs), cytokines, and downstream signaling pathways ^2^. Inhibition of this adaptive response by lapatinib prevents downstream Src, FAK, and Akt signaling ^3^. Mechanistically, kinase inhibitor treatment can decrease shedding of RTKs, thus enhancing bypass signaling ^4^. Drug treatment can also cause changes in the cell cycle, promoting resistance in the quiescent cells adapting to the treatment ^5^. These examples demonstrate that quick and widespread changes in signaling can occur in response to anti-cancer drugs.

The tumor microenvironment is a complex, heterocellular system with known roles in modulating tumor progression and drug response ^6^. The extracellular matrix (ECM) is one feature of the microenvironment that can alter cell signaling and facilitate adaptive resistance. For example, ECM stiffness can activate RTKs and engage integrin binding to promote cell growth and survival through the Raf/MEK/ERK and PI3K/Akt/mTOR pathways ^7,8^. Our group and others have employed engineered biomaterials to elucidate drug resistance in response to biophysical and biochemical stimuli from the ECM. Two-dimensional (2D) hydrogels allow for the presentation of controlled integrin binding, a physiological stiffness, and are amenable to drug screening. Three-dimensional (3D) hydrogels also present physiological dimensionality and biochemistry through matrix degradability, matrix adhesion, and homotypic or heterotypic cell-cell interactions. No single system can capture all features of the tumor microenvironment, and complex systems do not allow for isolation of the effect of individual cues. Here, we propose a biomaterial-based method to systematically vary stiffness, dimensionality, and cell-cell contacts to analyze matrix-mediated adaptive resistance by measuring genetic and phospho-signaling changes.

All of these extracellular cues can alter the signaling networks targeted by cancer therapeutics. For example, β^1^ integrin-mediated tolerance to a BRAF inhibitor *in vivo* can be recapitulated by stiff synthetic matrices *in vitro* ^9^. We recently developed a high-throughput drug screening platform based on a poly(ethylene glycol)-phosphorylcholine (PEG-PC) hydrogel and identified that increased substrate stiffness imparts resistance to sorafenib ^10,11^. Similarly, others have demonstrated decreased drug sensitivity in 3D hydrogels compared to 2D hydrogels and tissue culture polystyrene (TCPS) ^12^. These studies demonstrate the vast opportunities synthetic biomaterials provide for exploring drug resistance and rewiring mediated by features of the ECM but have yet to capitalize on the potential for a systematic analysis of stiffness, geometry, and cell-cell contacts afforded by such systems to understand mechanisms of ECM-mediated resistance.

It is imperative to consider multiple components of the ECM while studying drug response, as variations in matrix complexity can produce considerably different predictions for *in vivo* drug response to anti-cancer drugs. Since no single system can capture all features of the tumor ECM, we investigated drug resistance and adaptive reprogramming of breast cancer cells across multiple biomaterial microenvironments to analyze the genetic and phospho-signaling contributions to adaptive resistance. Multiple linear regression modeling revealed that MEK signaling explained the difference in RTK-targeted drug response across the ECMs, and co-administering a MEK inhibitor with sorafenib improved efficacy of the single agent *in vitro* and *in vivo*. This is the first report to combine systems biology analysis with screening across biomaterial platforms to identify MEK as a mediator of ECM-driven resistance to RTK inhibitors.

## Materials and Methods

### Cell culture

All supplies were purchased from Thermo Fisher Scientific (Waltham, MA) unless otherwise noted. Human breast cancer cell lines MDA-MB-231 and SkBr3 cells were generous gifts from Dr. Sallie Smith Schneider at the Pioneer Valley Life Sciences Institute and Dr. Shannon Hughes at the Massachusetts Institute of Technology, respectively, and were cultured in Dulbecco’s Modified Eagle’s Medium (DMEM) supplemented with 10% fetal bovine serum (FBS) and 1% penicillin-streptomycin (P/S) at 37ºC and 5% CO_2_.

### Polymerization of 2D and 3D PEG hydrogels

2D PEG-phosphorylcholine (PEG-PC) hydrogels were prepared and protein was coupled as described previously ^10^. Glass-bottom 96-well plates (no. 1.5 coverslip glass; In Vitro Scientific, Sunnyvale, CA) were plasma treated (Harrick Plasma, Ithaca, NY) and subsequently methacrylate-silanized with 2 vol % 3-(trimethoxysilyl) propyl methacrylate (Sigma-Aldrich, St. Louis, MO) in 95% ethanol (pH 5.0) for 5 min, washed 3 times with 100% ethanol, and dried at 40°C for 30 minutes. PEGDMA (Mn 750, Sigma-Aldrich), is added at 1.1 wt % (10 kPa, 0.015 M) or 3.2 wt % (33 kPa, 0.043 M) to a solution of 17 wt % 2-methacryloyloxyethyl phosphorylcholine (PC) (Sigma-Aldrich) in phosphate-buffered saline (PBS). Solutions were sterilized with a 0.2 μm syringe filter (Thermo) and degassed by nitrogen sparging for 30 seconds. Free-radical polymerization was induced by addition of 0.05 wt % ammonium persulfate (APS, Bio-Rad Laboratories, Hercules, CA) and 0.125 vol % tetramethylethylenediamine (TEMED, Sigma-Aldrich). Hydrogels of 40 μL per well in the 96-well plates were polymerized under nitrogen for 30 minutes. Post-polymerization, hydrogels were allowed to swell for 24 hours in PBS, then treated with 100 μL of sulfo-SANPAH (ProteoChem, Denver, CO; 0.6 mg/mL in pH 8.5 HEPES buffer) under UV light for 20 minutes, rinsed twice with HEPES buffer, and followed immediately by incubation with type I collagen at 3.3 μg/cm^2^ (Thermo) overnight.

3D PEG-maleimide (PEG-MAL) hydrogels were prepared at 10 wt % (3 kPa) and 20 wt % (5 kPa) solution with a 20K 4-arm PEG-MAL (Jenkem Technology, Plano, TX) and 2mM of the cell binding peptide CRGD (Genscript, Piscataway, NJ), and was crosslinked at a 1:1 ratio with 1K linear PEG-dithiol (Sigma-Aldrich) in 2mM triethanolamine (pH ~ 7.4) ^13^. Hydrogels were polymerized in 10 μL volumes for 5 minutes before swelling in cell culture medium.

Bulk diffusion of Rhodamine 6G (R6G) (Stokes’ radius 0.76 nm, Sigma-Aldrich) in 3D PEG-MAL hydrogels was measured by encapsulating 0.1 g/L R6G in the hydrogel and sampling the supernatant at 5-minute intervals. The samples were analyzed on a fluorescent plate reader at excitation: 526 nm/emission: 555nm. The diffusion coefficient of R6G was calculated using the Stokes’-Einstein equation.

### Formation of uniform tumor spheroids

Lyophilized poly(N-isopropylacryamide)-poly(ethylene glycol) (polyNIPAAM, Cosmo Bio USA, Carlsbad, CA) was reconstituted in cell culture medium according to the manufacturer’s instructions and kept at 4ºC until use. Cells were suspended in polyNIPAAM solution placed on ice at a density of 100,000 cells/mL for SkBr3 cells and 167,000 cells/mL for MDA-MB-231 cells, and each gel was made at a volume of 150 μL. Gelation occurred after 5 minutes at 37ºC, and gels were swollen in cell culture medium. Single cells were grown into spheroids for 14 days in the gels with regular medium changes. Spheroids were either dissociated back onto TCPS (for RNA-Seq and drug dosing) or encapsulated in 3D PEG-MAL gels (for RNA-Seq, drug dosing, and signaling) for 24 hours.

### Hydrogel mechanical characterization

Mechanical compression testing for 2D and 3D hydrogels was performed with a TA Instruments AR-2000 rheometer (New Castle, DE) at a 2 μm/s strain rate and analysis as previously described ^11^. Indentation testing for polyNIPAAM hydrogels was performed using a custom-built instrument previously described ^14^. For this application, a flat punch probe with a diameter of 1.5 mm was applied to samples at a fixed displacement rate of 5 μm/s, for maximum displacement of 100 μm. PolyNIPAAM samples were placed on a heated surface to maintain a sample temperature between 37-40**°**C throughout testing.

### Quantification of drug response

At 24 hours post-seeding, cells on biomaterials were treated with lapatinib (0-64 μM, LC Laboratories, Woburn, MA), sorafenib (0-64 μM, LC Laboratories), temsirolimus (0-80 μM, Selleckchem, Houston, TX), doxorubicin (0-32 μM, LC Laboratories), or dimethyl sulfoxide (DMSO, Sigma-Aldrich) as a vehicle control. Where noted, inhibitors were included in the medium: PD0325901 (3 μM, Sigma-Aldrich), SP600125 (20 μM, LC Laboratories). After 48 hours of drug treatment, cell viability was quantified by CellTiter Glo Luminescent Cell Viability Assay (Promega, Madison, WI), and luminescence was read on a Biotek Synergy H1 plate reader after incubation for 10 minutes. Separately, at 24 hours, viability of untreated cells was determined via CellTiter Glo for a measurement of initial seeding density. IC_50_ and GR_50_^15^ values were calculated using the initial seeding density and drug treated measurements using GraphPad Prism v6.0h.

### Cell seeding on biomaterials

Cells were seeded on and in biomaterial microenvironments in serum-free DMEM (1% P/S, 20 ng/mL of epidermal growth factor (EGF), R&D Systems, and 20 ng/mL of platelet-derived growth factor BB (PDGF-BB), R&D Systems, Minneapolis, MN) or serum containing DMEM (1% P/S and 10% FBS) where noted. Cells were seeded at 30,000 cells/cm^2^ on PEG-PC gels and on tissue culture plastic (TCPS), or as single cells at 500 cells/μL in 3D PEG gels. Spheroids were recovered via addition of cold serum free medium (1% P/S) on ice for 5 minutes for gel dissolution, and then gravity sedimentation for 30 minutes on ice. The supernatant was removed and the spheroids collected from each polyNIPAAM gel were re-suspended in PEG-MAL solution to make 9 3D hydrogels.

### RNAseq

Total RNA was isolated using Gen Elute mammalian total RNA miniprep kit (Sigma-Aldrich). The Illumina TRUseq RNA kit was used to purify and fragment the mRNA, convert it to cDNA, and barcode and amplify the strands. Quality and length of the inserts was confirmed with an Agilent Genomics 2100 bioanalyzer, followed by single-end reads on a NextSeq 500 to generate a complete transcriptome from each sample. Transcripts were aligned to the hg19 human reference genome using the Tuxedo Suite pathway ^16^.

### Gene Set Enrichment Analysis

Aligned samples were converted to GCT and CLS file formats using useGalaxy.org ^17^. Samples were separated by material condition across both cell lines to determine biomaterial microenvironment-specific gene categories. GSEA software was used to analyze enriched KEGG pathways as previously described ^18^, and pathways with significant enrichment of expressed genes were selected ^19^.

### Multiplex phospho-protein quantification

Cells were seeded onto 2D hydrogels at 56,600 cells/cm^2^, or into 3D gels at 200,000 single cells in 20 μl volume gel. Spheroids were grown for 14 days in polyNIPAAM, and then transferred to 3D PEG-MAL gels, where spheroids from three polyNIPAAM gels were transferred to four 20 μl 3D PEG-MAL gels. Samples were seeded in serum free DMEM. After 24 hours, cells were dosed with either 8 μm sorafenib, 8 μm lapatinib, 25 μm temsirolimus, or DMSO control in serum free medium (or serum containing medium where noted) for 4 hours, then washed once with ice cold PBS and lysed in RIPA buffer supplemented with protease (EDTA-free protease inhibitor cocktail tablets, 1 tablet in 10 ml, Roche, Indianapolis, IN) and phosphatase (1x phosphatase inhibitors cocktail-II, Boston Bioproducts, Boston, MA) inhibitors, 1 mM phenylmethylsulfonyl fluoride (Thermo Fisher Scientific), 5 μg/ml pepstatin A (Thermo Fisher Scientific), 10 μg/ml of leupeptin (Thermo Fisher Scientific), 1 mM sodium pyrophosphate (Thermo Fisher Scientific), 25 mM β-glycerophosphate (Santa Cruz, Dallas, TX). Protein concentration was determined with a BCA assay (Sigma-Aldrich). PEG-MAL gels were manually dissociated with a pipet tip for 30 seconds after addition of lysis buffer.

Lysate concentrations were adjusted to 520 μg/ml and then analyzed with the MILLIPLEX MAP Multi-Pathway Magnetic Bead 9-Plex - Cell Signaling Multiplex Assay (Millipore, Billerica, MA; analytes: CREB (pS133), ERK (pT185/pY187), NFκB (pS536), JNK (pT183/pY185), p38 (pT180/pY182), p70 S6K (pT412), STAT3 (pS727), STAT5A/B (pY694/699), Akt (pS473) supplemented with additional EGFR (pan Tyr) and MEK1 (pS222) beads according to the manufacturer’s instructions, with the exception that beads and detection antibodies were diluted four-fold. Controls were performed to ensure that samples were within the linear dynamic range. Data is reported as the Mean Fluorescent Intensity (MFI) or fold change in MFI relative to the vehicle control.

SkBr3 cells on TCPS, 33 kPa 2D gels, and spheroids in 3 kPa PEG-MAL were serum starved for 24 h and then stimulated with 100 ng/mL of EGF (R&D Systems, Minneapolis, MN), and cell lysates were collected at 0, 5, 15, 60 minutes, and 24 hours time points. Lysate concentrations were adjusted to 160 μg/mL, and samples were quantified the MILLIPLEX MAP Phospho Mitogenesis RTK Magnetic Bead 7-Plex Kit (Millipore; analytes: c-Met/HGFR (panTyr), EGFR (panTyr), ErbB2/HER2 (panTyr), ErbB3/HER3 (panTyr), ErbB4/HER4 (panTyr), IR (panTyr), and IGF1R (panTyr) according to the manufacturer’s instructions, with the exception that beads and detection antibodies were diluted four-fold.

### Experimental Model And Subject Details

Multiple linear regression modeling was performed using’stats::lm’ within R, regressing each compendium of signaling measurements against the corresponding viability measured in the same conditions. All relationships were assumed to be additive, and no interaction or intercept terms were included in the models. Each measurement was z-score normalized before regression. The two-sided p-values presented were calculated based upon the significance of each parameter being non-zero. The overall performance of each model (as calculated by the F-statistic) corresponded well to the significance of individual terms. All code and raw measurements for model development are available on Github (<u>https://github.com/meyer-lab/Barney-Peyton</u>).

### Immunofluorescence and immunohistochemistry

Cells were grown in serum for 24 h in the indicated material condition, fixed, and stained for Ki67. Cells were serum-starved and stimulated with EGF as described above, and stained according to standard protocols for total EGFR or HER2. The following antibodies were used for immunofluorescence: Ki67 (ab16667, 1:200, Abcam), EGFR (D38B1, 1:100, Cell Signaling Technology), and HER2 (29D8, 1:200, Cell Signaling Technology). All tumor samples were fixed and paraffin embedded. Ki67 staining (ab16667, 1:100, Abcam) was done on 6 μm tissue slices. All Alexa Fluor secondary antibodies (Thermo Fisher Scientific) were used at 1:500. Samples were imaged on a Zeiss Cell Observer SD (Carl Zeiss AG, Oberkochen, Germany).

### Animal Xenografts

All animal experiments were in accordance with institutional guidelines and approved by the Institutional Animal Care and Use Committee at the University of Massachusetts Amherst. UMass has an approved Animal Welfare Assurance (#A3551-01) on file with the NIH Office of Laboratory Animal Welfare. MDA-MB-231 cells were suspended in Matrigel (Corning, Corning, NY), and 10^6^ cells were subcutaneously injected into the mammary fat pad of 8-10 week old female NOD scid gamma mice from a breeding colony. Each mouse was injected with two tumors. All mice were housed in ventilated cages with sterile bedding, food, and water. Injected cells were grown *in vivo* for 7 days then drugs were suspended in DMSO and administered with an intraperitoneal (IP) injection daily for 14 days using a 27-gauge needle. Mice received 100 μL of one of five different treatments: vehicle (100% DMSO), sorafenib at 10 mg/kg, PD0325901 at 10 mg/kg, or a combination of the drugs at 5 or 10 mg/kg each. A minimum of 4 mice were analyzed for each drug dosing condition, and each tumor was considered one replicate.

### Quantification And Statistical Analysis

Prism v5.04 (GraphPad Software) was used to perform unpaired Student’s t-test, a one-way analysis of variance (ANOVA) with a Tukey post-test. A two-way ANOVA determined how the IC_50_ changed with respect to material modulus, geometry, and medium condition, with a Bonferroni post-test (GraphPad Prism v5.04). A two-way ANOVA, performed in R, was used to determine drug contribution to tumor burden. Data are reported as mean ± standard error, where p ≤ 0.05 is denoted with *, ≤ 0.01 with **, and ≤ 0.001 with ***.

## Results

### Geometry impacts innate breast cancer cell response to RTK/MAPK-targeting drugs

Distinct differences across *in vitro* screening methods, including stiffness, dimensionality, and cell-cell interactions prevent comparisons across existing studies. Therefore, we developed a system to independently evaluate the effects of both the geometry and stiffness of the microenvironment on breast cancer cell response to targeted and non-targeted drugs. We measured the GR_50_ (concentration of drug which dampens growth by 50%) ^15^ for four drugs (sorafenib, lapatinib, temsirolimus, and doxorubicin) across multiple biomaterials: TCPS, a 2D PEG-PC hydrogel, and a 3D PEG-MAL hydrogel with encapsulated single cells or tumor spheroids (Figure 1a). To create tumor spheroids, single cells were encapsulated sparsely in polyNIPAAM hydrogels and grown into uniform clonal spheroids over 14 days in culture (Figure 1b-d). This 14 day endpoint achieves a relatively homogeneous population of viable multicellular tumor spheroids with an average diameter less than 100 μm (Figure 1b-c) ^20^. This diameter was chosen to ensure drug ^21^ and oxygen diffusion into the spheroids. Spheroids were transferred to PEG-MAL hydrogels for dosing, where bulk diffusion measurements of rhodamine 6G suggest small molecules diffuse through the 3D hydrogel at 2.5x10^−6^ cm^2^/s (data not shown), which means that the drug will reach the cells within 10 seconds. Further, immunofluorescent staining of Ki67 expression revealed that there were proliferating cells throughout the entire spheroid, suggesting there were no significant nutrient or oxygen diffusion limitations within the 3D PEG hydrogel (Figure S1).

**Figure 1.**
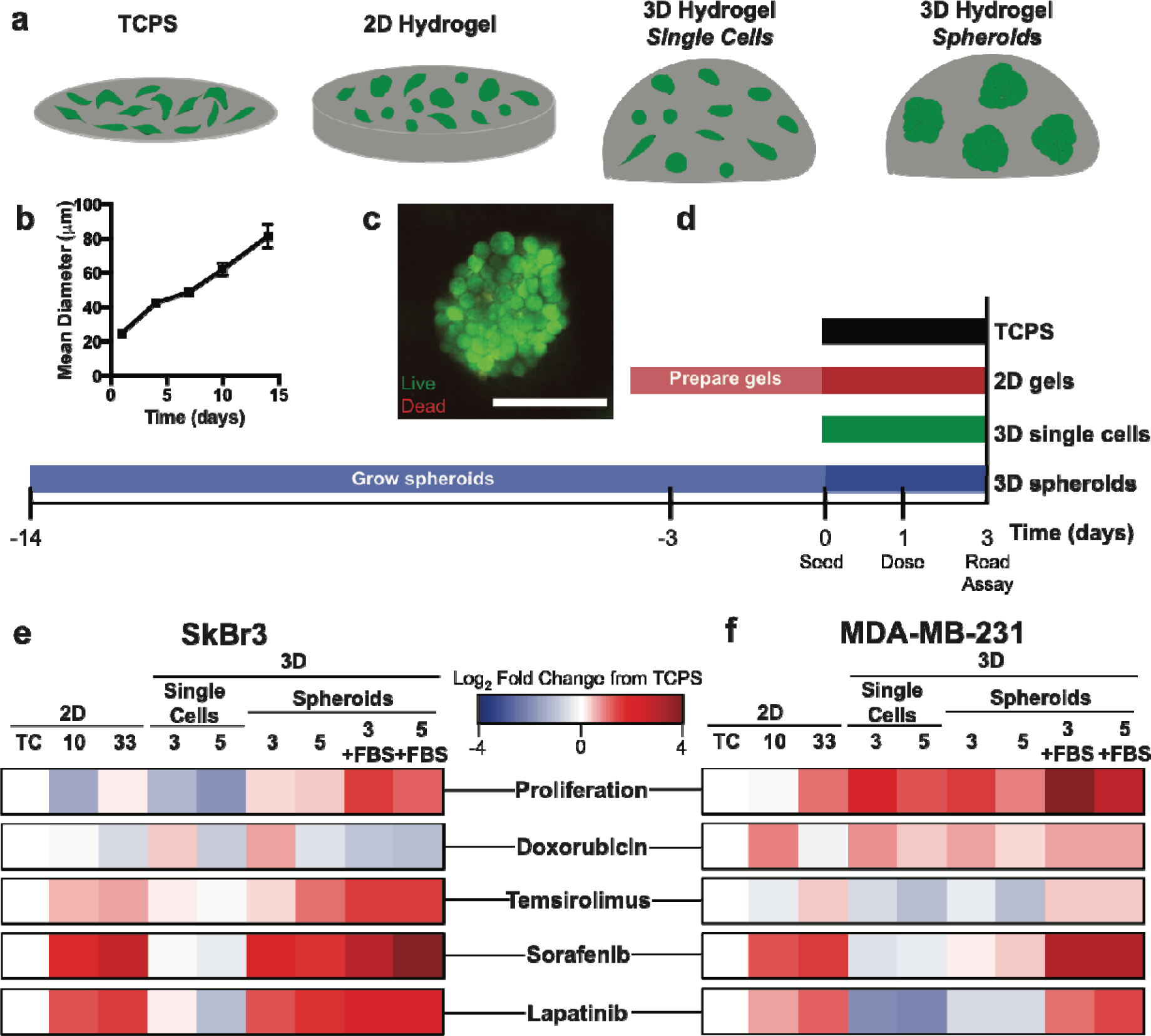
Drug response is dependent upon the biomaterial screening platform. a. Schematic of biomaterial drug screening platforms, including TCPS, 2D hydrogels linked with collagen I, and 3D hydrogels with RGD for cell adhesion, either with encapsulated single cells or tumor spheroids. Each hydrogel condition was screened at two different moduli (2D: 10 or 33 kPa, 3D: 3 or 5 kPa). b. Quantification of mean spheroid diameter for MDA-MB-231 spheroid growth in polyNIPAAM gels over 14 days. c. Viability of MDA-MB-231 spheroid after 14 days of growth in polyNIPAAM, and encapsulation in 3 kPa 3D PEG-MAL hydrogel for 3 days. Green: live cells, red: dead cells. Scale: 100 μm. d. Experimental time investment required for the different screening platforms. Timeline displays preparation time required for each system prior to treating with varying concentrations of drug. e-f. SkBr3 (e) and MDA-MB-231 (f) proliferation after 3 days of culture (top row), and GR_50_ in response to doxorubicin, temsirolimus, sorafenib, an lapatinib across the different biomaterial platforms. Red: increase from TCPS control (resistance compared to TCPS), blue: decrease from TCPS control (sensitivity compared to TCPS). TC: TCPS, 10: 10 kPa 2D hydrogel, 33: 33 kPa 2D hydrogel, 3: 3 kPa 3D hydrogel, 5: 5 kPa 3D hydrogel, +FBS: cells were treated in 10% serum-containing medium. Unless noted by +FBS, all cells were treated with drugs in serum-free medium containing EGF and PDGF-BB.

We previously found that breast cancer cells on stiffer substrates were more resistant to sorafenib ^10^, so we included two different moduli for the 2D and 3D hydrogels (Figure S2). Although the effect of altered proliferation across the biomaterials (Figure 1e-f) would confound comparisons of traditional IC_50_ calculations, it is accounted for in the GR_50_ calculation ^15^. Therefore, the differences we report in drug response are a direct result of ECM-driven resistance, independent of a growth advantage. In Figure 1e-f, drug response is reported as a fold change in GR_50_ from the indicated condition compared to TCPS, where red indicates resistance and blue indicates sensitivity compared to TCPS. Response to doxorubicin did not vary significantly across geometry, modulus, or medium (with or without serum, varied for the spheroid model only) in either cell line (Figure 1e-f, Figure S3). Response to temsirolimus was not dependent on platform in the MDA-MB-231 cells, while in the SkBr3 cells, geometry and medium, but not modulus, had a small, but significant, impact on GR_50_. However, sorafenib and lapatinib response varied considerably across the biomaterial platforms, particularly in the MDA-MB-231 cells (Figure 1e-f, Figure S3). It is notable that the SkBr3 response to sorafenib was only dependent on geometry, and not the addition of serum, suggesting a unique role for the ECM in mediating drug response, independent of exogenous mitogenic factors. A two-way analysis of variance (ANOVA) showed that the changes in geometry and medium had a larger effect on total variance than the change in modulus as reported by the percent of variation across biomaterial platforms (Table S1). Within each geometry, we observed no significant difference in drug response at either stiffness, and we encourage the reader to view the bar graphs and statistical analysis provided in the electronic supplementary information (Table S1, Figure S3), indicating that cancer cell resistance to sorafenib and lapatinib was most sensitive to changes in geometry within this modest stiffness range.

### Gene expression is not responsible for ECM-driven differences in drug response

We performed RNA-seq on MDA-MB-231 and SkBr3 cells cultured on TCPS, on 2D hydrogels, as tumor spheroids, and as a tumor xenograft (Figure 2a). Hierarchical clustering of differentially expressed genes first separated samples by the cell line, then by material platform (Figure S4). The biomaterial platform dictated clustering within each cell line and dimensionality had greater influence on clustering than stiffness. Gene expression on the 2D hydrogels (10 and 33 kPa) was similar for both cell lines, and these samples are closely related to cells on TCPS. Toward understanding whether this was an immediate effect of signals from the ECM or from selective growth of a subset of cells in different environments, we also analyzed cells initially grown into spheroids, but then dissociated and cultured back on TCPS for 24 hours. The gene expression profile from these dissociated spheroids was also closely related to the other 2D conditions, indicating this expression was likely derived from the assay platform, rather than the culture platform (Figure 2a).

**Figure 2.**
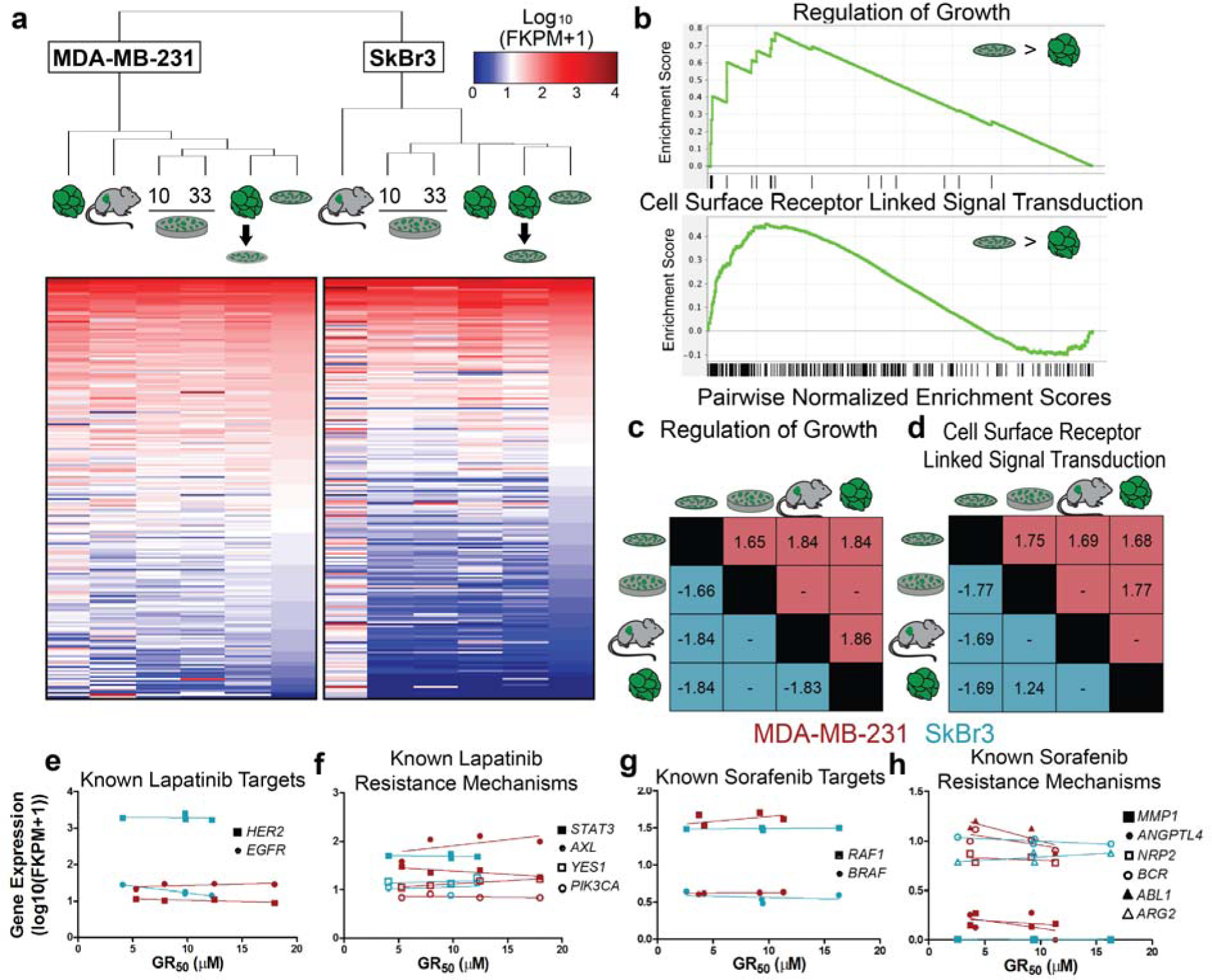
Expression of growth and signal transduction genes distinguishes screening platforms but does not explain drug response. a. Clustering and normalized expression values of significantly differentially expressed genes across the biomaterial platforms for MDA-MB-231 and SkBr3 cell lines. Genes were ordered by expression on TCPS, and each cell line was independently filtered for the top 250 genes by maximum expression across all platforms. All significantly differentially expressed genes (p<0.01) are shown in Figure S4. MDA-MB-231 order of platforms: spheroids, xenograft, 2D hydrogels at 10 kPa, and 33 kPa, spheroids reseeded to TCPS, TCPS. SkBr3 order of platforms: xenograft, 2D hydrogels at 10 kPa, and 33 kPa, spheroids, spheroids reseeded to TCPS, TCPS. Blue: low expression, red: high expression b. Representative gene set enrichment curves for regulation of growth (MDA-MB-231, TCPS over 3D spheroids) and cell surface receptor linked signal transduction (MDA-MB-231, TCPS over 3D spheroids). c-d. Normalized enrichment scores in pairwise platform comparisons for (c) regulation of growth (d) and cell surface receptor linked signal transduction. MDA-MB-231 comparisons are enriched in red and SkBr3 comparisons are in blue. Comparisons are shown as *ROW* condition over *COLUMN* (i.e., Top row shows MDA-MB-231 enrichment on TCPS over other platforms). – indicates no data. e-h. Expression of known targets for (e) lapatinib and (g) sorafenib and previously published off-target genes for (f) lapatinib and (h) sorafenib across the biomaterial platforms with linear correlations.

Gene set enrichment analysis (GSEA) identified several significantly enriched pathways in one or more of the biomaterials. For example, genes associated with the regulation of growth and receptor-linked signal transduction are upregulated in MDA-MB-231 cells on plastic compared to MDA-MB-231 cells in 3D spheroids (Figure 2b). While this is not an exhaustive list, cells on TCPS expressed more cytokine and associated receptor genes and cells in the xenograft upregulated expression of ECM components, particularly collagens and keratins. In Figure 2c and d, the normalized enrichment scores are plotted as pairwise comparisons, enriched in the row condition over the column platform in red for MDA-MB-231 cells and in blue for SkBr3 cells. As expected, both cell lines upregulated genes associated with growth regulation on TCPS compared to all the other conditions we examined (Figure 1e-f, Figure 2b-c). Furthermore, the MDA-MB-231 xenograft tumors upregulated growth-related genes compared to cells grown *in vitro* as spheroids, suggesting that the mouse host environment provided additional growth cues (Figure 2c). However, both geometry and stiffness influenced surface receptor linked signal transduction. For example, cells on TCPS universally showed enrichment for these genes compared to the other biomaterials, and our spheroids showed enrichment over the 2D hydrogels in the SkBr3 cells (Figure 2d).

We then used this expression data to determine if ECM-mediated drug response was a result of changes in the gene expression of drug targets for lapatinib and sorafenib across the biomaterials (Figure 1e-f). We found no correlation between expression of these target genes and the GR_50_ (Figure 2e,g). However, both of these drugs are known to alter signaling of many other pathways, and a literature search identified several known unintentional targets or resistance mechanisms for each drug ^22-26^ (Figure 2f,h). Although many of these genes showed little to no change in expression across the biomaterials, we did note that *AXL* expression showed a slight positive trend, but not a significant correlation, with lapatinib response in the MDA-MB-231 cells (Figure 2f). AXL has previously been shown to predict drug resistance to ErbB targeted inhibitors in many cancer types ^26^, but does not appear to significantly contribute to the drug responses examined here.

### Multiple linear regression reveals that using a MEK inhibitor with sorafenib is an effective combination therapy

Since there were large differences in GR_50_ across the screening microenvironments, and known drug targets were not responsible for ECM-mediated resistance, we hypothesized that plasticity in intracellular signaling networks was responsible for the differential responses. We measured a panel of phospho-proteins associated with RTK signaling at basal levels and in response to drug treatment across the biomaterial ECMs. Here, we chose to closely examine sorafenib and lapatinib across material platforms, and added temsirolimus to compare to a drug whose response was largely independent of ECM (Figure 1e-f).

The basal levels of nearly every analyte were significantly lower in 3D than on 2D ECMs (Figure 3a-b), which matches growth and signal transduction gene signatures from our expression data (Figure 2c-d). As expected, drug treatment reduced the phosphorylation level of the intended targets (Figure 3c-d). However, we only observed significant reduction in known targets and increases in adaptive pathways in 2D, suggesting that the 3D environment suppresses drug efficacy through dampened signaling and kinome rewiring. Dimensionality-dependent signaling differences were also present during direct activation of the receptor by its cognate ligand. When stimulated with epidermal growth factor (EGF) on TCPS, pEGFR was elevated within 5 minutes, and EGFR was internalized within an hour (Figure S5a-c). When spheroids were encapsulated in 3D hydrogels, pEGFR and pHER2 were still stimulated at 5 minutes, but only modestly, and EGFR localization did not change with EGF stimulation (Figure S5a-f).

**Figure 3.**
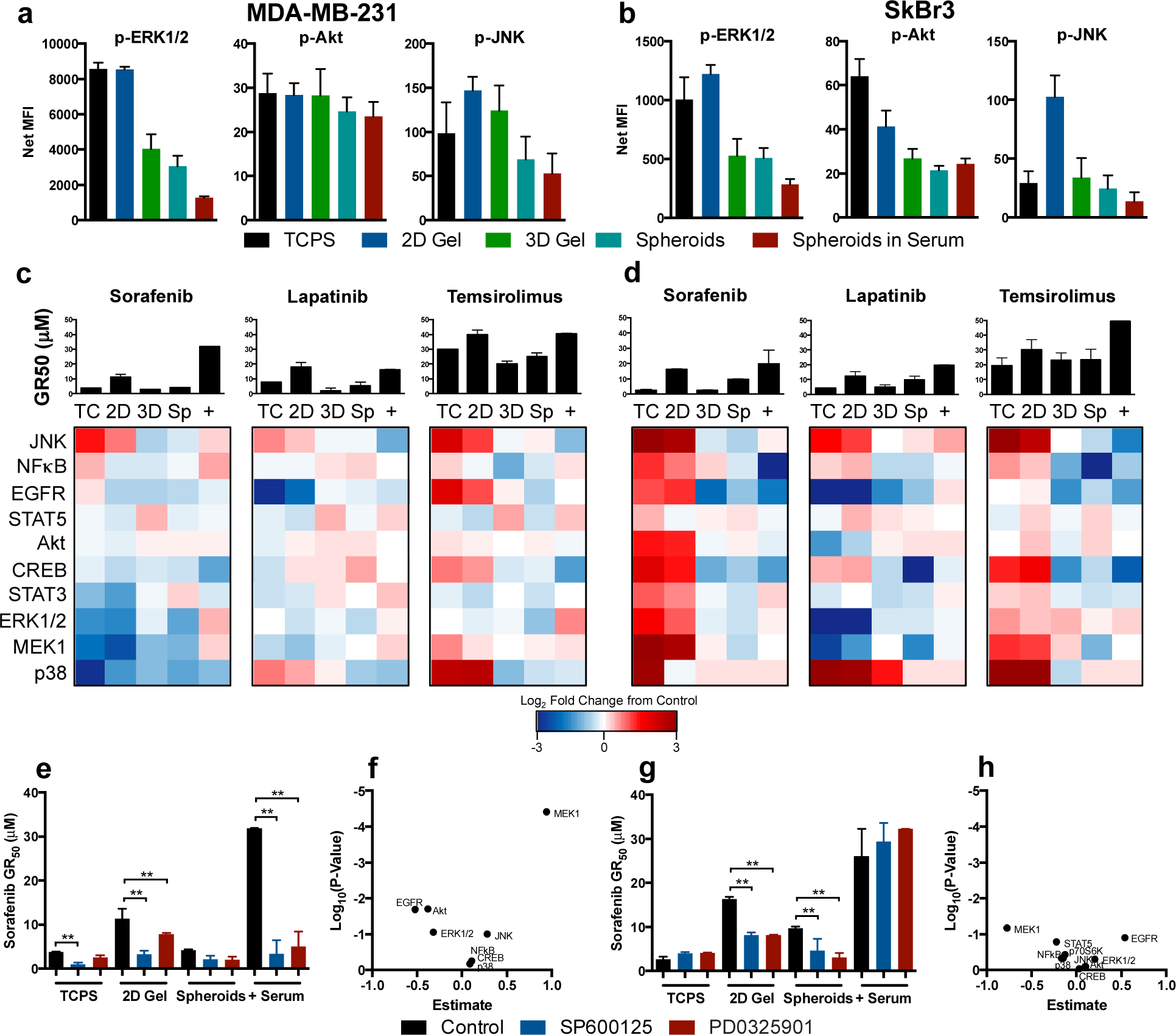
Systems analysis reveals that MEK regulates ECM-mediated sorafenib efficacy. a-b. Basal p-ERK (pT185/pY187), p-Akt (pS473), and p-JNK (pT183/pY185) signaling for MDA-MB-231 (a) and SkBr3 (b) cell lines across the five screening microenvironments. Black: TCPS, dark blue: 2D gel, green: 3D gel with single cells, teal: 3D gel with spheroids, red: 3D gel with spheroids in serum. 2D hydrogel: 33 kPa, 3D hydrogel: 3 kPa. c-d. Phospho-signaling in response to drug treatment and GR_50_ for MDA-MB-231 (c) and SkBr3 (d) cell lines. TC: tissue culture polystyrene, 2D: 2D PEG-PC hydrogel, 3D: single cells encapsulated in 3D hydrogels, Sp: spheroids encapsulated in 3D hydrogels, +: spheroids in 3D hydrogels in serum containing medium. Phospho-proteins measured include: CREB (pS133), ERK (pT185/pY187), NFκB (pS536), JNK (pT183/pY185), p38 (pT180/pY182), STAT3 (pS727), STAT5A/B (pY694/699), Akt (pS473), EGFR (pan Tyr), and MEK1 (pS222). Red: increase compared to vehicle control, blue: decrease from vehicle control. e,g. Sorafenib GR_50_ alone or in combination with SP600125 (JNK inhibitor, 20 μM) or PD0325901 (MEK inhibitor, 3 μM) for MDA-MB-231 (e) and (g) SkBr3 cell lines. Black: control, blue: SP600125, red: PD0325901. f,h. MLR models of signaling data for MDA-MB-231 (f) and SkBr3 (h).

Our lab has previously demonstrated the role of JNK in stiffness-dependent sorafenib resistance ^10^. Because we observed increased pJNK in response to the drug treatments in both cell lines, we hypothesized that JNK may mediate drug response across the biomaterials. Co-administration of JNK inhibitor SP600125 sensitized the MDA-MB-231 cells to sorafenib on 2D biomaterials and as spheroids with serum, as well as TCPS (Figure 3e). SP600125 only sensitized the SkBr3 cells in the spheroid and 2D hydrogel conditions, but not the stiff TCPS environment, perhaps a result of the very high increase in pJNK in the sorafenib-treated SkBr3 cells in that environment (Figure 3g). We separately examined JNK phosphorylation after treatment with doxorubicin, a non-targeted drug that provoked low variability across platforms, and found that JNK signaling was still altered by drug treatment. This suggests that here JNK may be caused by drug-induced stress, but does not significantly contribute ECM-mediated resistance (Figure S6a-b). Further, the efficacy of the JNK inhibitor in the MDA-MB-231 spheroids in the 3D hydrogel was mirrored in the simple TCPS assay, and did not need screening across biomaterials for discovery. Thus, we sought to identify an efficacious combination therapy using this multi-environment screening approach that could not be identified through simple TCPS screening.

Multiple linear regression (MLR) models revealed non-intuitive relationships between the sorafenib-, lapatinib-, and temsirolimus-treated signaling and drug response data for both cell lines across the biomaterials. This model was constructed from phosphorylation data of 10 analytes 4 hours after targeted drug treatment at 15 conditions, including 3 drugs, and 5 biomaterial platforms per drug. Although targeting the MEK/ERK pathway was not an obvious choice from the signaling data (Figure 3c-d), these models revealed that variability in MEK phosphorylation was the key factor associated with viability in response to drug treatment in MDA-MB-231 cells (Figure 3f,h). Here it is important to note that the MLR output includes the contribution of each signaling analyte to drug response, and that while MEK was identified as the key factor, other analytes have non-zero contributions to the model. To determine the necessity of modeling cell response across several biomaterial platforms and drugs, we performed subsequent analyses where we independently excluded either all 3D conditions (single cells, spheroids, spheroids with serum) or 2 of the drugs (lapatinib and temsirolimus). Neither of these modified models achieved significance (Figure S7a). Returning to our model that includes all 3 drugs in the MDA-MB-231 cells, we determined that each 3D condition could be excluded independently, but that TCPS, 2D biomaterials, and at least one 3D condition were necessary to predict MEK phosphorylation as a likely mechanism of ECM-mediated resistance (Figure S7b).

We next co-administered both sorafenib and the MEK inhibitor PD0325901 to cells in the different biomaterials. In both cell lines, there was no improved efficacy when co-treating sorafenib and PD0325901 on TCPS (Figure 3e,g). However, both cell lines were sensitized to sorafenib when co-administered with PD0325901 in the 2D hydrogel and spheroid conditions (Figure 3e,g). This suggests MEK may be a more physiological resistance mechanism to sorafenib, as SkBr3 cells did not show significant variability of MEK across platforms, but were still responsive to co-treatment in our biomaterials. Interestingly, the MDA-MB-231 cells are KRAS mutant, while the SkBr3 cells are not, meaning this treatment is independent of both this mutation and their clinical subtype. Altogether, the identification of MEK as an effective target for combination therapy was only realized through screening across many biomaterials and using a systems approach to analyze a phospho-signaling dataset. Likewise, the efficacy of the MEK inhibitor in combination with sorafenib was only achieved through screening in the more complex 2D or 3D biomaterial microenvironments, and would be missed using TCPS or via genetic analysis alone.

**Figure 4.**
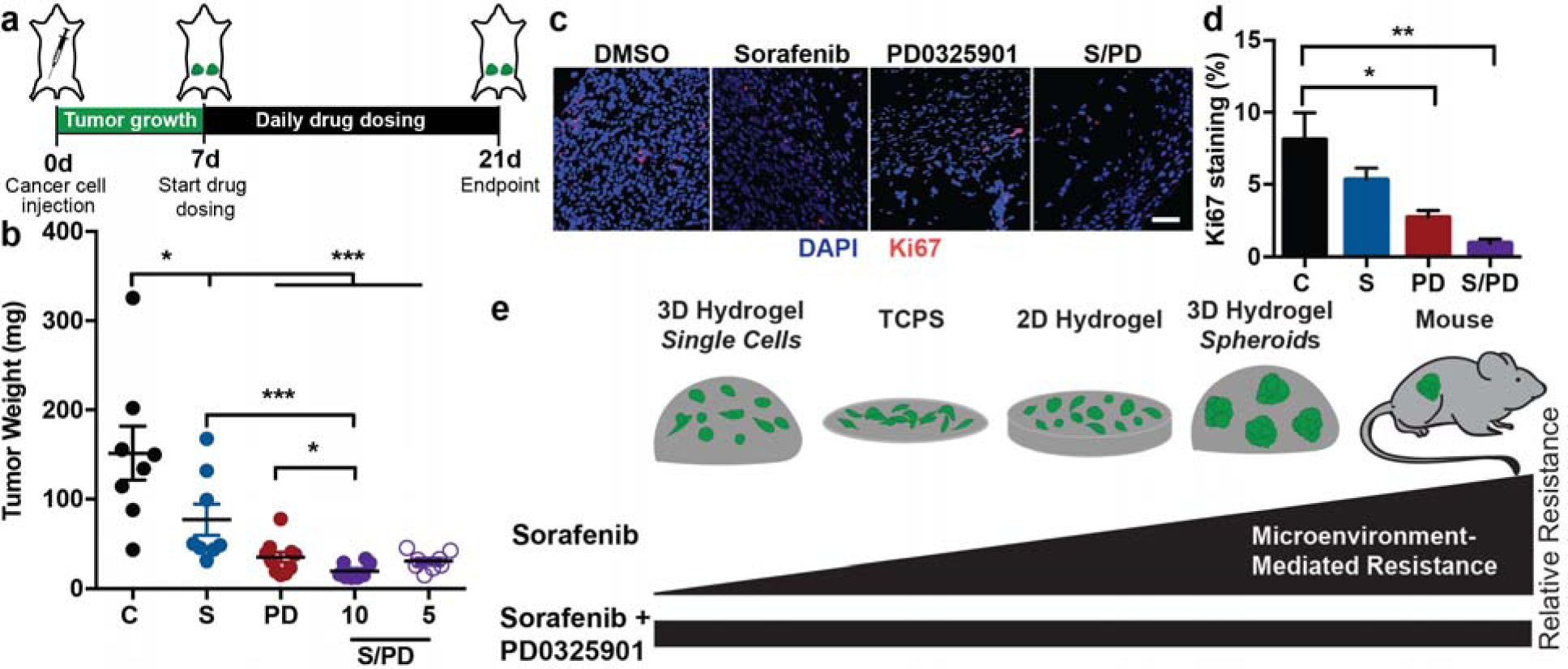
PD0325901 and sorafenib are an effective combination therapy to reduce *in vivo* tumor burden. a. Schematic for tumor induction of the MDA-MB-231 cells, grown first for 7 days, followed by daily drug dosing for 14 days until a 21-day endpoint. b. Final tumor weight for each drug dosing condition (C: control, S: 10 mg/kg sorafenib, PD: 10 mg/kg PD0325901, and co-dosing at either 5 mg/kg or 10 mg/kg of each therapeutic, p = 0.05; two-tailed t-test on log transformed data). c. Representative images for Ki67 staining in tumors and d. quantification of Ki67 positive cells in drug treatment groups (S/PD is the 10 mg/kg combination therapy). Scale bar is 50 μm. e. Screening for ECM-mediated resistance in drugs like sorafenib can be used in combination with multiple linear regression modeling to find combination therapies with the MEK inhibitor PD0325901. Error bars represent mean and SEM, N≥4.

### MEK inhibition improves sorafenib efficacy *in vivo*

Given the promising *in vitro* results with the MEK inhibitior, we tested PD0325901 as a combination therapy with sorafenib in an *in vivo* model. MDA-MB-231 tumors were inoculated in the mammary fat pad of NSG mice and allowed to grow for 7 days, at which point all mice had palpable tumors. From days 7 to 21, we treated daily with one of four regimens: 1) vehicle control (DMSO), 2) 10 mg/kg sorafenib, 3) 10 mg/kg PD0325901, or 4) co-treatment of 10 mg/kg each sorafenib and PD0325901 (Figure 4a). As combination therapies are often associated with increased adverse events ^27^, we also included a fifth treatment group which received a lower dose of the combination therapy at 5 mg/kg each of sorafenib and PD0325901. All of these doses were found to be well tolerated by mice in separate experiments (data not shown), although future work will need to better optimize the doses of this combination therapy for optimal clinical efficacy. PD0325901 was the most effective single agent inhibitor and, as predicted in the 2D hydrogel and spheroid screening models, co-treating mice with PD0325901 and sorafenib had the most significant reduction in tumor burden, cellularity, and proliferation (Figure 4b-d). In the lower concentration combination therapy (5 mg/kg each), reduction in tumor weight was comparable to 10 mg/kg PD0325901 alone (Figure 4b). A two-way ANOVA showed that PD0325901 (F_value_=54.7, p<0.001) contributed more to the decrease in tumor burden than sorafenib (F_value_=8.2, p<0.01). The combination therapy of sorafenib and PD0325901 additively decreased tumor burden (F_value_=0.3, p=0.6).

## Discussion

By screening drug response across many ECMs, we identified that breast cancer cell response to sorafenib (Ras/Raf/MEK/ERK targeted) and lapatinib (EGFR-HER2 targeted) was significantly dependent upon the biomaterial environment (Figure 1e-f). Gene expression of known drug targets was unchanged across the ECMs (Figure 2e-h), but a systems analysis of phospho-kinome signaling in response to drug treatment across our different biomaterials revealed MEK as a key contributor to the variation in observed ECM-mediated resistance (Figure 3f). Although the combination therapy did not improve drug response in cells on TCPS, a MEK inhibitor in combination with sorafenib eliminated the matrix-mediated response in biomaterials *in vitro* and reduced tumor burden *in vivo* (Figure 4e).

While most current literature examines signaling changes within a single platform ^28-31^, we applied MLR across five ECM conditions and three targeted drug treatments to determine which phospho-proteins varied significantly with ECM-mediated drug response. This approach captures multiple aspects of the ECM that may contribute to drug response *in vivo*, including ECM stiffness, dimensionality, and cell-cell contacts using both 2D and 3D biomaterials. Synthetic 2D biomaterials allow for the presentation of full length proteins, while spheroids encapsulated in 3D better recapitulate tumor geometry. Importantly, cells seeded onto 2D biomaterials or as single cells in 3D only resided in that platform for 24 hours, so they likely relied exclusively on the provided collagen I (2D) or RGD (3D) for biochemical signals. However, breast cancer cells grown as spheroids upregulate ECM genes, such as fibronectin, suggesting additional ECM cues might contribute to resistance ^20^. Recent work has shown that the ECM-mediated signaling can impart resistance to combined inhibition of PI3K and HER2 ^32^, but our method identified that targeting two nodes in the same pathway, Raf with sorafenib and MEK with PD0325901, is a promising combination therapy for breast cancer. This non-intuitive co-treatment, which would not be identified by screening on traditional TCPS, suggests a similar approach may be beneficial in identifying combination therapies for kinome-targeting drugs, where adaptive resistance is frequently mediated by the ECM ^10, 12^.

Combination therapies with MEK inhibition are effective in many cancer types ^33-36^. In fact, MEK inhibition is currently under evaluation in clinical trials because many cancers activate MEK signaling as a form of acquired resistance ^37^. Such combination therapies are often designed to limit adaptive survival signaling, and generally target two distinct pathways. Interestingly, in our work, co-targeting MEK improved sorafenib efficacy *in vitro* and in a pre-clinical *in vivo* tumor regression model, even though both drugs target the same pathway (Figure 4b). Others have seen similar improved efficacy in HER2+ breast cancer when co-treating the HER2-targeted drugs lapatinib and trastuzumab, suggesting that combination therapies targeting the same pathway can have improved effects by overcoming acquired resistance ^38^. However, not all cells within a tumor contribute equally to drug response. Tumors are highly heterogeneous and subpopulations can evade both cytotoxic and targeted therapeutics via upregulation of drug transporters, mutations of drug targets, adaptive signaling, and other mechanisms ^2,39,40^. Here we identified the additive effect of co-dosing with sorafenib and a MEK inhibitor on bulk tumor properties, but further work is necessary to understand the contribution of individual populations to resistance.

We measured a large panel of phospho-proteins to see if the biomaterial environment facilitated phospho-signaling changes in response to the drug treatments that were associated with drug sensitivity or resistance. In our soft 3D environments, basal and drug-stimulated signaling were significantly suppressed, in agreement with other data in spheroids and on soft substrates ^41-43^. JNK appears to be a broad adaptive response ^44,45^; however, MLR identified MEK as the most significant driver of ECM-mediated resistance to sorafenib. ERK is downstream of both the lapatinib and sorafenib targets, which may explain why MEK/ERK signaling was so critical to drug response in our data. In agreement with our findings, myeloid leukemia cells have a stiffness-dependent response to drugs, including sorafenib and a MEK inhibitor, which target the RAF/MAPK pathway ^46,47^. We have previously shown that sorafenib resistance increased with increasing hydrogel stiffness ^10^, and many others have also demonstrated that stiffness is a significant driving force for drug resistance ^9,10,28,29,48-51^. Thus, we were surprised to find that drug resistance was only minimally affected by stiffness within the modest range examined here (Figure 1e-f). However, in our data, dimensionality contributed more to drug resistance than stiffness. Previous work has similarly shown that cancer cell spheroids are more resistant to paclitaxel than a monolayer ^30^, and the matrix-attached cells in spheroids are resistant to PI3K/mTOR inhibition ^31^.

Because cells are less able to form cell-matrix adhesions on soft versus stiff 2D substrates, and in 3D versus 2D environments ^52^, genes associated with integrin-mediated adhesion and cell-surface receptor linked signal transduction are varied across our biomaterials (Figure 2b-d). Matrix adhesion via integrins mediates survival and drug resistance through activation of survival signaling, such as FAK, Src, and ERK ^53-55^. Disruption of cell adhesion can sensitize tumor cells to drug treatments ^10,32,56-58^. In 3D, we generally observed greater drug sensitivity for single cells compared to spheroids (Figure 1e-f), perhaps because single cells have lower overall adhesions to surrounding cells and matrix ^12^. Spheroids and tumors can also exhibit multicellular drug resistance through either cell-cell contact-mediated signaling or drug diffusion limitations ^59^, and often co-targeting multiple pathways is required to sensitize the outer, matrix-attached cells ^31^. While our spheroid size was intentionally kept small to minimize diffusion limitations, the cell-cell contact inherent to the spheroid structure may provide survival signaling, resulting in increased resistance to the RTK-targeted drugs.

In our hands, gene expression of drug targets was relatively unchanged across our materials. Previous work has reported significant changes in gene expression profiles of cell lines cultured in 2D and 3D laminin-rich ECM ^60^. This is similar to our own data, where we observed that surface receptor linked signal transduction genes vary with both geometry and stiffness, and are universally upregulated during culture on TCPS compared to the other biomaterials (Figure 2b). Also, several ligand-receptor pairs were co-expressed and enhanced *in vivo*, including BMP and IL6 signaling (Figure S8a-b). We found that growth context determined the expression of cell surface receptors, while expression of the associated ligands was dependent on the current assay platform and was more sensitive to stiffness.

Interestingly, both gene expression and drug sensitivity were regained when cells grown as spheroids were dissociated and seeded back onto TCPS (Figure 2a, Figure S8). This demonstrates that drug response in the timeframe we examined is dependent on the current environment and not necessarily a result of stable genetic alterations or subpopulations resulting from a previous growth condition. In our gene expression data, the xenograft was the most distant from other samples for the SkBr3 cells, but not for the MDA-MB-231 cells. This is likely due to the differences in length of time required for the xenografts to reach a similar size. The MDA-MB-231 xenograft tumor reached 50 mm^3^ within 10 days, and the SkBr3 xenograft took 3 weeks to reach 50 mm^3^. In an *in vivo* microenvironment, xenograft tumors derived from highly lung metastatic clones of the MDA-MB-231 breast cancer cell line have a distinct ECM signature that up-regulates growth factor signaling including TGF-β and VEGF ^61^. We conclude that although gene expression may be useful to inform patient treatment ^62^, it was not sufficient to predict *in vitro* or pre-clinical drug efficacy here.

## Conclusions

Though the ECM can impart resistance to some RTK inhibitors (sorafenib and lapatinib), MLR modeling of phospho-signaling identified combinatorial strategies to combat this resistance (Figure 4e). This approach could provide insight into other anti-cancer drugs with varied clinical success, such as neratinib ^63,64^ and sunitinib ^65,66^. However, not all the drugs we tested showed ECM-dependent responses, and response to a cytotoxic drug, such as doxorubicin, would have been predicted in a simple, quick, cost-effective TCPS assay. It is therefore critical to weigh the monetary cost and time burden of the screening approach against the potential benefits. The efficacy of the PD0325901 and sorafenib combination therapy would not have been realized using only one screening environment or without a systems biology analysis. We envision this approach can be used to identify drugs with poor *in vivo* efficacy early in the drug development pipeline and identify novel drug combinations to overcome ECM-induced resistance.

## Acknowledgements

We thank Shannon Hughes and Sallie Smith Schneider for generously providing cell lines and Jungwoo Lee for providing the mice. We acknowledge the University of Massachusetts Genomics Resource Laboratory for their assistance with sequencing, and our use of the gene set enrichment analysis (GSEA) software, and Molecular Signature Database (MSigDB). This work was funded by an NSF-NCI award DMR-1234852, a grant from the NIH (1DP2CA186573-01) and a CAREER grant from the NSF (1454806) awarded to S.R. Peyton. S.R. Peyton is a Pew Biomedical Scholar supported by the Pew Charitable Trusts. L.E. Barney was partially supported by National Research Service Award T32 GM008515 from the National Institutes of Health. A.D. Schwartz was supported by a National Science Foundation Graduate Research Fellowship (1451512). T.V. Nguyen and S.R. Peyton were supported by a Barry and Afsaneh Siadat Career Development Award. A.S. Meyer was supported by an Early Independence Award from the NIH (DP5-OD019815).

